# Idiosyncratic Tower of Babel: Individual differences in word meaning representation increase along abstractness

**DOI:** 10.1101/2020.08.28.272997

**Authors:** Xiaosha Wang, Yanchao Bi

## Abstract

Humans primarily rely on language to communicate, based on a shared understanding of the basic building blocks of communication: words. However, words also have idiosyncratic aspects of meaning. Do we mean the same things when we use the same words? Classical philosophers disagreed on this point, speculating that words have more similar meanings across individuals if they are either more experiential (John Locke) or more abstract (Bertrand Russell). Here, we empirically characterize the individual variation pattern of 90 words using both behavioral and neuroimaging measures. We show that the magnitude of individual meaning disagreement is a function of how much language or sensory experience a word associates with, and this variation increases with abstractness of a word. Uncovering the cognitive and neural origins of word meaning disagreements across individuals has implications for potential mechanisms to modulate such disagreements.

---

Language is a proxy for human minds. We transfer our thoughts across individuals and across time and space through language. We often assume that differences in thoughts are reflected by different choices of words, and that speakers of the same language have common conceptual understandings about the basic elements of word meaning. Such commonality is the basis of effective learning and communication when speaking the same language and word meaning misalignment is usually only discussed within the context of cross-language speakers ^1,2^. However, there are also intriguing individual variations in how we understand a word within a language, which associate with important nonverbal properties, such as political position ^3^ or emotional perception ^4^. The question of individual variation in word representation has intrigued classical philosophers including John Locke and Bertrand Russell, who put forward opposite speculations: that words denoting “complex ideas” (e.g., abstract words) may have lower inter-subject consistency (ISC) ^5^; and that words entailing more “abstractness of logic” may have greater individual consistency ^6^. Here we empirically quantify, using both behavioral and neural signatures, the consistency and variations of word meaning representations across speakers of the same language and from a relatively homogeneous culture and education group, and investigate the underlying mechanisms leading to individual variation.

The nature of and variables affecting individual variation of word meaning are intrinsically related to the general principles of how word meanings are represented in the human brain. Meaningful variance in a system stems from the dimensions that make up the corresponding representation. Decades of research have focused on the common cognitive and neural basis of conceptual/semantic representations, converging on the consensus that they are compositional, entailing salient sensory, motor, and emotion-related attributes, and distributed over multiple systems of the cortex ^7–15^, with recent evidence for additional non-sensory, language-derived representations ^16,17^. We thus based our mechanistic investigation on these underlying attribute dimensions.

Experimental stimuli included 90 words covering key semantic domains whose cognitive and brain mechanisms have been extensively studied (**Table S1**), including those having external sensory referents (varying in sensory and motor-related attributes: animals, face/body parts, manmade objects) and those without specific external referents (non-object abstract words, including those with and without emotional associations, e.g., “violence” vs. “result”). We quantified their representations in each participant (all Chinese college students in Beijing) based on both behavioral judgments (Experiment 1) and brain activation patterns measured by functional neuroimaging (Experiment 2). We computed the ISC for each word from behavioral (ISC-behavior) and brain (ISC-brain) data, and examined the effects of key representational dimensions in predicting these ISC values.

## Cognitive representations of word meaning: Individual consistency predicted by Language/Sensory Experiences

We constructed cognitive word meaning representations from behavioral judgments of the semantic distances among 90 words. This approach was taken because word meaning representation is difficult to capture by explicit definitions ^18,19^ and it is assumed to be (at least partly) represented by the relationships with other words ^3,4,20–22^. We asked 20 subjects (college students from Beijing) to rate meaning distance among 90 words using a multi-arrangement method ^23^, **Fig. 1a**), which produced a 90 × 90 representational dissimilarity matrix for each subject. We visualized these distance matrices using multidimensional scaling in a 2-D plot; words spatially closer are more semantically related (**Fig.1b** and **Fig. S1a**). The mean Fisher-Z-transformed Pearson correlations of the entire 90-word matrices between each subject and the group (in a leave-one-subject-out fashion) are 0.58 ± 0.10, indicating medium-level consistency across subjects (**Fig. S1b** and **S1c**). Next we computed the word-level cognitive inter-subject consistency. For each word, we took the rated distances with all other words, i.e., an 89-dimension vector, as the “representation” of this word for each subject. Then we computed the Pearson R between each subject pair, Fisher-Z-transformed, and averaged across all subject pairs, giving the ISC-behavior of this word.

**Fig. 1.**
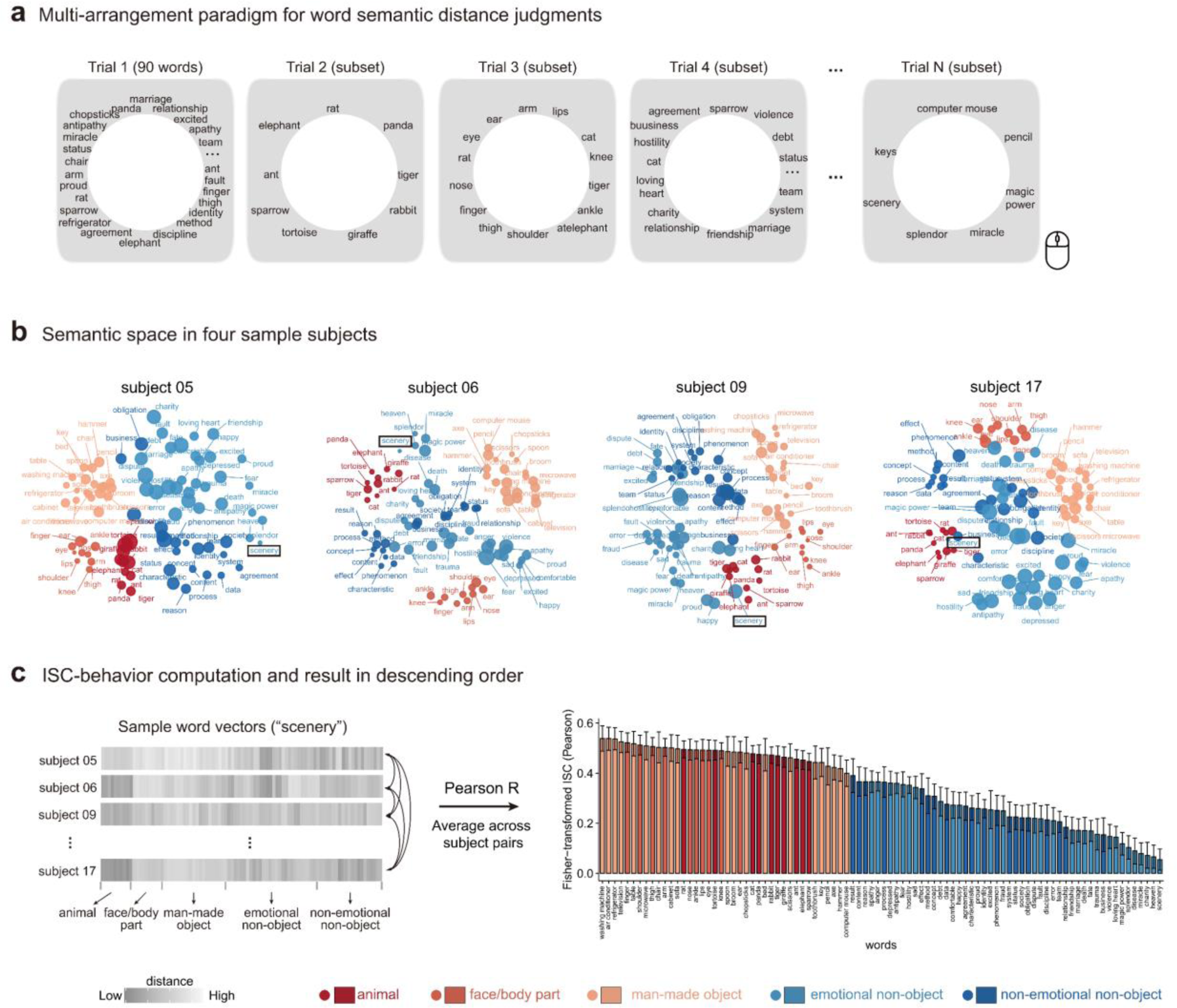
Inter-subject consistency of word meanings based on behavioral judgments. **(a)** We adopted the multi-arrangement paradigm to evaluate the semantic representations of 90 words from five semantic domains ^23^. In this task, subjects were asked to drag and drop words in a circular array on a computer screen according to their semantic distances. The first trial presented all 90 words and yielded a 90 × 90 matrix containing Euclidean distances among all the words. The subsequent trials presented adaptively-selected subsets of words that had been clustered together in previous trials, producing partial distance matrices. The task lasted for 60 min and the output was a 90 × 90 distance matrix of weighted average distances across the multiple arrangements. **(b)** Visualization of semantic space using multidimensional scaling (MDS) of individual semantic distance matrices in 4 sample subjects (the smacof package in R, type = interval; see **Fig.S1a** for other subjects’ semantic spaces). Dot sizes reflect stress values, with larger dots indicating higher stress (i.e., greater inconsistency between original space and MDS space). **(c)** Each word for each subject was represented as an 89-dimension vector reflecting its semantic associations with other words. Pearson correlation coefficients were computed for each pair of subjects, then Fisher-Z-transformed and averaged across subject pairs to obtain ISC-behavior. The error bars of ISCs were calculated as the standard deviations of the bootstrap distribution of 10,000 resamplings of subjects (see **Fig. S2** for error bars generated by bootstrap resampling of words).

As evident from the bar plots in **Fig. 1c** (and **Fig.S2**), words referring to concrete referents (objects) such as *washing machine* and *finger* have systematically and significantly higher ISC-behavior than words that do not refer to specific external referents (e.g., “business”, “scenery”; mean Fisher Z*-*transformed R: 0.48 ± 0.03 vs. 0.24 ± 0.09; independent-samples *t*_*(88)*_ = 15.81, *p* = 1.93 × 10^−27^). That is, on average, people’s meaning representations for words with specific sensory referents are about twice as similar as those for non-external referent (abstract) words.

Next, we examined the mechanistic origins of word meaning variation across individuals: What aspects of word meaning representations account for the individual variation? Motivated by cognitive and neural theories, the following meaning dimensions were considered: external-referent related, including sensory experience (across all sensory modalities) and motor action experiences (manipulation, navigation, and stress-related actions) ^7,9,24,25^; emotion-related ^14^, including emotional valence and arousal; and language-related ^16^, i.e., language descriptivity. We asked independent groups of subjects (from the same linguistic/cultural background as the main experiments) to rate the 90 words on each dimension on a 7-point scale (see Methods and Materials for details). We computed the mean and variation (indexed by standard deviation, SD) (**Fig. 3a**) for each word across subjects’ ratings as candidate sources for the ISC-behavior.

**Fig. 2.**
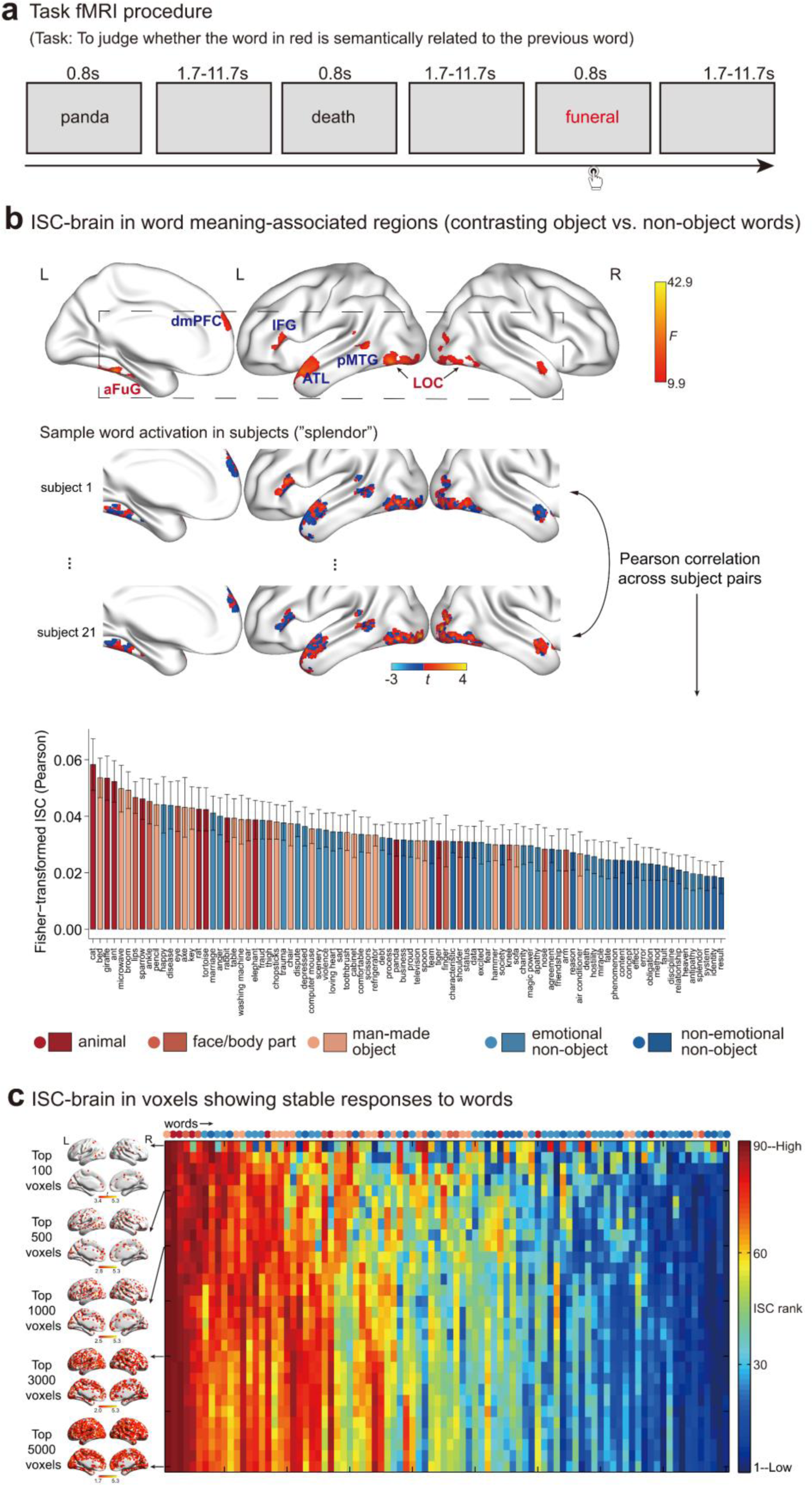
Inter-subject consistency of word meanings based on brain activation patterns. **(a)** Brain responses to each of 90 words were collected in a task fMRI experiment in which subjects were asked to think about word meanings and judge whether occasional words displayed in red (catch trials) are semantically related to the previous word. **(b)** ISC-brain in word meaning-associated brain regions, defined as regions whose activation strengths significantly differentiated between object and non-object words across 21 subjects (voxelwise *p* < .005, FWE-corrected cluster-level *p* < .05; clusters with blue labels are relatively non-object preferring, showing higher activations to non-object than object words; clusters with red labels are relatively object-preferring). For each word, the activation pattern in these regions in each subject was taken as its brain representation and Pearson correlation was computed for each pair of subjects, which were then Fisher-Z-transformed and averaged across subject pairs to obtain ISC-brain. The error bars of ISCs for each word were calculated as the standard deviations of the bootstrap distribution of 10,000 resamplings of subjects. **(c)** ISC-brain calculated from activation patterns in grey matter voxels showing consistently high stability to words across subjects. The brain images (visualized using BrainNet ^32^ with the “Maximum Voxel” algorithm) show the distribution of grey matter voxels with top-N highest stability scores (see Methods and Materials for details). The heatmap shows the ranking of ISC-brain values across 90 words in top N voxels we sampled (from top 100 to top 1000 voxels, in a step of 100 voxels, and from top 1000 to top 5000 voxels, in a step of 200 voxels). Words were sorted in descending order according to the averaged ISC-brain values from top 100 to 5000 voxels. LOC, lateral occipital cortex; aFuG, anterior fusiform gyrus; pMTG, posterior middle temporal gyrus; ATL, anterior temporal lobe; IFG, inferior frontal gyrus; dmPFC, dorsomedial prefrontal cortex. L, left hemisphere; R, right hemisphere.

**Fig. 3.**
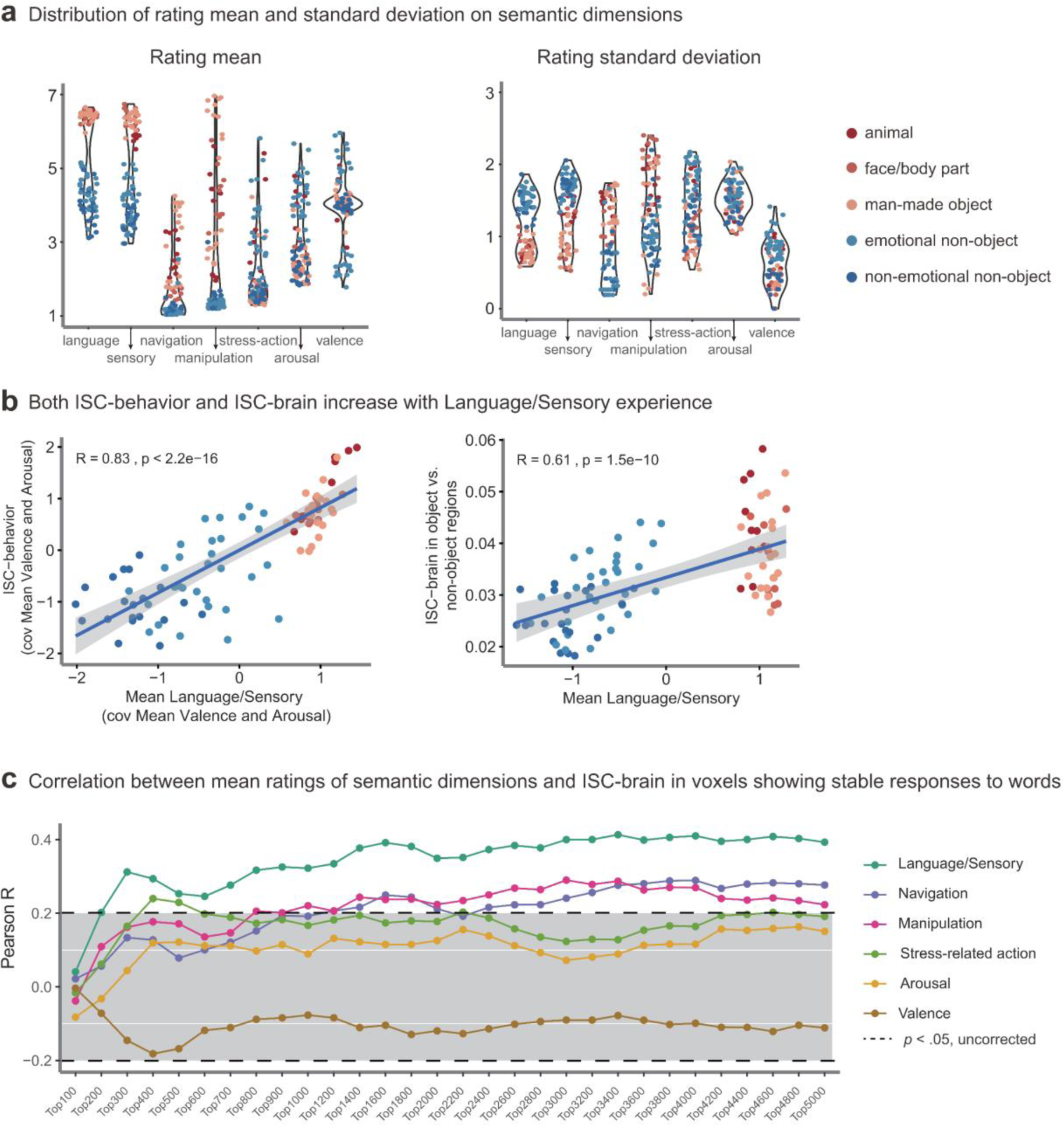
Cognitive mechanistic origins of ISC-behavior and ISC-brain. **(a)** Distributions of 90 words on each of seven semantic dimensions gleaned from cognitive and neuroscience studies. For each semantic dimension, ratings were collected from independent groups of 24-28 subjects (from the same linguistic/cultural background as the main experiments) on a 1-7 scale. Both rating mean and standard deviations (SD) across subjects were calculated as the candidate sources for individual variations. Language and sensory rating means were transformed into z-scores and averaged to obtain the Mean Language/Sensory variable due to high correlation between them. **(b)** Both ISC-behavior and ISC-brain increases with Mean Language/Sensory experiences. For ISC-behavior, the partial regression plot between ISC-behavior and Mean Language/Sensory experiences is shown here (the stepwise regression model identified the words’ mean ratings on Language/Sensory, Arousal, and Emotional Valence as significant predictors of ISC-behavior). For ISC-brain, Mean Language/Sensory was the only significant predictor in the regression model including significant predictors of rating mean and SDs. **(c)** Correlations between rating means of semantic dimensions and ISC-brain in voxels showing consistently high stability to words across subjects (sampled from top 100 to 5000 voxels) confirm the positive association of ISC-brain with Mean Language/Sensory experiences. The correlations with mean Navigation/Manipulation ratings were driven by their inter-correlations with Language/Sensory ratings revealed by partial correlation analyses: the effects of Mean Language/Sensory experiences still hold when controlling for Navigation or Manipulation ratings (*p*s < .034, for top 200 to 5000 voxels), and the effects of Navigation/Manipulation are no longer significant when controlling for Mean Language/Sensory experiences (*p*s > .17).

Each word’s ISC-behavior was predicted using multiple linear regression models with these variables as predictors. The means of language descriptivity and sensory experience are highly correlated across the 90 words (*r* = .94) and were collapsed by taking the averaged *z* values into a single Mean Language/Sensory variable (**Fig. S3**). The significant mean predictors (Mean Language/Sensory, Mean Arousal, and Mean Valence) and SD predictors (SD Language, SD Manipulation, and SD Valence) were obtained separately first, and then considered together (**Table S2**). The Mean Language/Sensory, Mean Arousal, and Mean Valence predictors are significant in the final model, together explaining 76.2% of the variance in the ISC-behavior: ISC-behavior = 0.74 × Mean Language/Sensory – 0.33 × Mean Arousal – 0.17 × Mean Valence + 0.59 (stepwise regression model; *F(3,86)* = 91.64, *p* = 1.07 × 10^−26^; see **Fig. 3b** for the partial regression plot between Mean Language/Sensory and ISC-behavior). These effects persist when including word frequency and familiarity as nuisance variables (**Table S2**). As an alternative approach to dealing with the correlated variables, we employed principal component analysis (PCA, **Table S3**), and results converged on the findings that the PC with high loadings of mean language-descriptivity and sensory-experience ratings was a significant predictor for ISC-behavior, and revealed that the PC composed of SDs of emotion-related variables was another significant predictor (see Supplementary Results and **Table S4**).

Note that in this way of computing word representation vectors, each word has its own set of reference points (the other 89 words). We performed validation analyses where the same set of reference points were used across words. We randomly split the 90 words into two 45-word sets, and for words in set 1 their rated distances to the 45 words in the other set were taken as the word-specific vector space. Across 10,000 random split-halves, the ISC-behavior values across words obtained in the two approaches (i.e., distance with own sets or common sets of words) are largely consistent: averaged Fisher z*-*transformed R (± SD) between the two approaches is 0.83 ± 0.30. The relationships between ISC-behavior and the above-mentioned significant predictors of semantic dimensions (i.e., Mean Language/Sensory, Mean Arousal, Mean Valence, SD Language, SD Manipulation, and SD Valence) were evaluated with a series of partial correlations (i.e., ISC-behavior was correlated with one semantic dimension while controlling for other dimensions). Partial correlation between ISC-behavior with Mean Language/Sensory remains significant (Fisher z*-*transformed R: 0.42 ± 0.19, 95% two-sided confidence interval (CI) based on percentile: 0.05 to 0.79, *p* = .024). The partial correlations between ISC-behavior and other semantic dimensions, especially emotion-related dimensions, are no longer significant when controlling for the other dimensions (Mean Arousal, Fisher Z*-*transformed R: −0.18 ± 0.15, 95% CI: −0.46 to 0.12, *p* = .21; Mean Valence, Fisher Z*-*transformed R: −0.27 ± 0.16, 95% CI: −0.58 to 0.06, *p* = .10; others: *p*s > .44). These results confirm that the more likely a word can be described using language and/or associates with sensory experience, the more similar semantic representations are found across people.

## Neural representations of word meanings: Individual consistency predicted by Language/Sensory Experiences

Words’ neural representations were constructed from fMRI BOLD signals. Twenty-one adult subjects, (20 of the same subjects from the multi-arrangement experiment), participated in an fMRI experiment. They read 90 words in the scanner (rapid event-related design, 10 repetitions for each word) and were asked to think about what the word meant and, when a word in red appeared (catch trials), to decide whether the word in red was semantically related to the previous word (**Fig. 2a**; see Methods and Materials for details). Brain activation patterns for each word in a mask comprising the word-meaning-associated regions were taken as its “neural representation”. Word-meaning associated regions were defined as clusters that were sensitive to major meaning type differences (contrasting objects vs. non-objects; voxelwise *p* < .005, FWE-corrected cluster-level *p* < .05; see below for alternative mask definitions s for validation). The group-level activation results include relatively object-preferring regions (bilateral lateral occipital cortex (LOC) and left anterior medial fusiform gyrus (aFuG)) and relatively non-object preferring regions (left posterior middle temporal gyrus (pMTG), bilateral anterior temporal lobes (ATL), left inferior frontal gyrus (IFG), and dorsal medial prefrontal cortex (dmPFC)) (**Fig. 2b**), which are highly consistent with the semantic literature ^17,26–28^. For each word, we obtained its activation pattern in this mask in each subject and calculated Pearson correlations of the activation patterns across all subject pairs, then Fisher-transformed and averaged the values to form the ISC-brain for each word.

As shown in the bar plots in **Fig. 2b**, words referring to concrete referents (objects) such as *cat* and *microwave* are again highly-significantly more consistent across individuals than words without external referents (mean Fisher z*-*transformed R: 0.039 ± 0.008 vs. 0.029 ± 0.007; independent-samples *t*_*(88)*_ = 6.23, *p* = 1.59 × 10^−8^). The ISC-brain and ISC-behavior is significantly correlated across words (*r* = .43, *p* = 2.20 × 10^−5^). Similar to the ISC-behavior analyses above, we examined which properties of word meanings account for the magnitude of ISC-brain across words. The Mean Language/Sensory is the only significant predictor in the final multiple regression model (**Table S5** and **Fig.3b**), explaining 37.4% of the variance in the ISC-brain: ISC-brain = 0.61 × Mean Language/Sensory + 0.033 (stepwise regression model, *F(1,88)* = 52.57, *p* = 1.53 × 10^−10^). The effect of Mean Language/Sensory persists when including psycholinguistic confounds (**Table S5**) and when using semantic PCs as predictors (Supplementary Results and **Table S4**). That is, the more likely a word can be described using language and/or associates with sensory experiences (typically those with an external referent), the more similar brain activation patterns it induces across individuals.

Validation analyses using four different word-related brain-mask definitions (see Methods and Materials for details) yielded largely-similar results to the analyses above. For Validation 1, without focusing on voxels showing different activations to pre-defined word types, we considered ISC in the whole brain, selecting grey-matter voxels showing consistently high stability to words across subjects (following ^21^, **Fig.2c** and **Fig.3c**). For Validation 2, we used the search term “word” in the meta-analysis platform Neurosynth ^29^ to identify brain areas consistently shown to be involved in word processing in the neuroimaging literature (**Fig.S4**). For Validation 3, in case any regions sensitive to word emotional meanings were not included in the main contrast above, we redefined the word-meaning-associated mask as those clusters sensitive to any differences among “object vs. emotional non-object vs. non-emotional non-object” (**Fig.S5**). For Validation 4, we calculated ISC-brain using voxels showing greatest sensitivity to object vs. non-object words in individual subjects (rather than the group) for each subject pair (within the group mask identified in the remaining 19 subjects, ^30^, **Fig.S6**). ISC-brain values obtained in these ways are highly correlated with the main results and all significantly predicted by Mean Language/Sensory experiences (**Fig. 2c, Fig.3c, Fig. S4-S6**).

Taken together, our results indicate that even speakers of the same language from a relatively homogeneous cultural/educational background exhibit substantial differences in what a word means, as measured by both behavioral ratings and brain activation patterns. In line with Locke’s (and not Russell’s) speculation, the individual consistency of meaning representations of words that designate concrete entities (i.e., with external referents) were significantly higher than those without external referents (abstract words), by both behavioral and brain measures. Also across both behavioral and brain measures, the magnitude of a word’s consistency across individuals was significantly positively predicted by the extent to which the word can be described using language and/or associates with sensory experiences. Further investigation is warranted about the origins of this variation, and whether and how culture, ideology, or contemporary artificial intelligence algorithms (i.e., automated individually tailored language and sensory inputs) may modulate individual differences through specific language and/or sensory experiences. What is clear is that human communication failures, especially in settings that rely largely on terms without external referents such as politics, sociology or legal domains, come from not only words’ sentential contexts but also the individual words themselves. Increasing language descriptiveness and sensory experiences may help reduce miscommunication originating on these basic elements, only on the basis of which more productive information exchanges and discussions are possible.

## Methods

### Participants

Twenty-one young healthy college students (11 females, mean age 21.1 years, ranging from 18 to 26 years) were recruited from university in Beijing for the study. All participated in the task fMRI experiment and 20 of them participated in the semantic distance judgment task. All participants were right-handed, native Chinese speakers, with at least one-year university studies in Beijing, and had normal or corrected-to-normal vision.

All participants provided informed consent and received monetary compensation for their participation. The study was approved by the Human Subject Review Committee at Peking University, China, in accordance with the Declaration of Helsinki.

### Stimuli

Stimuli in our study included 90 written words (Table S1), including 40 object words and 50 words without explicit external referents. Object words vary in their sensory and motor attributes, including 10 animals (e.g., *cat*), 10 face or body parts (e.g., *shoulder*), 20 man-made objects including tools and common household objects (e.g., *microwave*). Words without external referents vary in their emotional associations, containing 20 words without emotional connotations (e.g., *result*) (arbitrarily defined as having low arousal (<3) and being emotionally neutral (3.5-4.5) on 7-point ratings by independent groups of college students, see below) and 30 emotionally related words (e.g., *violence*). All words were highly familiar (mean ± SD = 6.5 ± 0.4, obtained from a 7-point familiarity rating on an independent group of 26 college students) and were disyllabic words except for five object words (in Chinese, “cat” and “bed” are monosyllabic; “giraffe”, “microwave”, and “washing machine” are tri-syllabic).

We compared different types of words on common psycholinguistic variables, including the number of strokes (a measure of visual complexity for Chinese words), word frequency, and subjectively rated familiarity. Compared with non-object words, objects had similar numbers of strokes (17.2 ± 5.8 vs. 16.1 ± 4.0; independent-samples *t*_*(88)*_ = 1.05, *p* = .30), were less frequent in a Mandarin Chinese corpus ^31^ (log word frequency: 1.0 ± 0.7 vs. 1.6 ± 0.7; *t*_*(88)*_ = −3.83, *p* < .001), but were rated more subjectively familiar (6.8 ± 0.2 vs. 6.2 ± 0.3; *t*_*(88)*_ = 12.0, *p* < .001). When further separating non-object words into emotional and non-emotional ones, we compared the three types of words on these variables using one-way analysis of variance (ANOVA), followed by Tukey *post hoc* test. The three types of words had similar numbers of strokes (*F(2,87)* = 1.84, *p* = .16). In word frequency, emotional non-object words (1.2 ± 0.5) were similarly frequent with object words (*p* = .42) and both were less frequent than non-emotional non-object words (2.2 ± 0.5; *p*s < .001). In subjectively rated word familiarity, emotional non-object words (6.2 ± 0.3) were similarly familiar with non-emotional non-object words (6.2 ± 0.2, *p* = .77) and both were rated less familiar than object words (*p*s < .001).

### Experiment 1: Word-level inter-subject correlation based on behavioral assessment

#### Semantic distance judgment task

The word meaning representations were obtained using a multi-arrangement paradigm ^23^. In this paradigm, subjects were asked to judge semantic distance among words by arranging them spatially close together or far apart in a circular array on a computer screen via mouse drag-and-drop operations (**Fig. 1a**). The task consisted of multiple trials: the first trial asked subjects to arrange all 90 words, producing an entire distance matrix, and the subsequent trials presented subjects with adaptively-selected word subsets that had been clustered together in previous trials, producing partial distance matrices. The task lasted for one hour, during which subjects completed various numbers of trials (range: 24-284; mean ± SD = 85 ± 71). The final distance measure for each subject is calculated as the weighted average of distance measures of their multiple arrangements.

#### Word-level ISC-behavior computation

To compute the word-level inter-subject consistency (ISC) in behavior, for each subject, each word was represented as an 89-dimension vector of its semantic distance with the remaining words. Pearson correlations among each pair of subjects; words were then computed, Fisher-Z-transformed, and then averaged across 190 subject pairs (20 subjects in total) to obtain ISC-behavior for each word. The confidence interval of ISC for each word was assessed in the following two approaches: 1) bootstrapping the subject set with replacement for 10,000 times, which evaluates ISC robustness across subjects; 2) bootstrapping the word set with replacement for 10,000 times, which evaluates ISC robustness across words included for judgment.

#### Validation of word ISC-behavior computation

Another way to construct word representations is based on their associations with a common set of base words. To compare the two approaches, we randomly split the 90 words into two word sets and represented each word in the first word set using its distance with the remaining 44 words in the same set or with the 45 words in the other set. ISCs were then computed from these data. This procedure was repeated for 10,000 times.

### Experiment 2: Word-level inter-subject correlation based on brain activation patterns

#### Task fMRI procedure

During the fMRI task, subjects were instructed to view each of 90 target words, think about their meanings, and perform an oddball one-back semantic judgment task, which was to judge whether occasional words in red were semantically related to the previous word by pressing the corresponding buttons with the right index or middle finger (catch trials). There were 10 runs (360 s per run). Each run consisted of 90 2.5 s-long word trials (0.8-s word followed by 1.7-s fixation), 14 2.5 s-long catch trials, and 30 2.5 s-long null trials, with the mean interval between two words being 3.23 s. Each target word appeared once within each run; the order of 90 target words was randomized for each run in each subject. Each run began with a 12s fixation period and ended with a 13 s rest period during which subjects received a verbal cue that the current run was about to end.

#### Image acquisition

All functional and structural MRI data were collected using a Siemens Prisma 3T Scanner with a 64-channel head-neck coil at the Center for MRI Research, Peking University. Functional data were acquired with a simultaneous multislice echoplanar imaging sequence supplied by Siemens (62 axial slices, repetition time (TR) = 2000 ms, echo time (TE) = 30 ms, multi-band factor = 2, flip angle (FA) = 90°, field of view (FOV) = 224 mm × 224 mm, matrix size = 112 × 112, slice thickness = 2 mm, gap = 0.2 mm, voxel size = 2 × 2 × 2.2 mm). A high-resolution 3D T1-weighted anatomical scan was acquired using the magnetization-prepared rapid acquisition gradient echo sequence (192 sagittal slices, TR = 2530 ms, TE = 2.98 ms, inversion time = 1100 ms, FA = 7°, FOV = 224 mm × 256 mm, matrix size = 224 × 256, interpolated to 448 × 512, slice thickness = 1 mm, voxel size = 0.5 × 0.5 × 1 mm).

#### Data preprocessing

Functional images were preprocessed using SPM12 (Wellcome Trust Center for Neuroimaging, London, UK, http://www.fil.ion.ucl.ac.uk/spm12/). For each individual participant, the first 4 volumes of each functional run were discarded for signal equilibrium. The remaining images were corrected for slice timing and head motion and spatially normalized to Montreal Neurological Institute (MNI) space via unified segmentation (resampling into 2 × 2 × 2 mm voxel size). No participant had head motion larger than 2 mm/2°. These images were directly submitted to general linear modeling (GLM) for multivariate pattern analyses and were further spatially smoothed using a 6 mm full-width half-maximum Gaussian kernel for univariate contrast analyses.

#### Computation of whole-brain activation patterns for each word

Whole-brain activation patterns for each word were obtained using a GLM with spatially normalized, unsmoothed functional images. For each subject, the GLM included for each run 90 regressors corresponding to the onsets of each target word and one regressor indicating catch trials, convolved with a canonical hemodynamic response function (HRF), and 6 head motion parameters. A high-pass filter cut-off was set as 128 s. The resulting *t* maps for each target word versus baseline were used to compute ISC-brain.

#### Word-level ISC-brain computation

The procedure for ISC-brain computation included the following steps: 1) Define word-associated voxels; 2) Extract activation patterns of each word from these voxels in each subject; 3) Compute, for each word, the Pearson correlations of activation patterns for each pair of subjects, which were Fisher-Z-transformed and averaged across subject pairs to obtain the ISC-brain of this word. The key step here was the definition of word-associated regions and we adopted the following approaches to validate the ISC-brain results. Functional activation maps were thresholded at voxelwise *p* < .005, FWE-corrected cluster-extent *p* < .05, unless stated explicitly otherwise.

First, we defined word-related regions as those sensitive to major meaning differences of objects vs non-objects. A GLM was built with spatially smoothed functional images and included regressors corresponding to the onsets of each word type (i.e., object and non-object) and one regressor for catch trials, together with 6 head motion parameters, for each run. The object vs. non-object contrast was computed in each subject and the resulting beta-weight images were submitted to an F-test at the group level.

Second, word-related regions were defined as grey matter voxels showing the most stable responses across words in 10 repetitions ^21^. For each of the voxels with probability higher than 0.4 in the SPM grey matter mask, we computed a stability score to evaluate its response consistency towards 90 words across 10 repetitions. In each subject, a grey matter voxel was assigned a 90 × 10 matrix, where the entry at row *i*, column *j*, was the beta weight of this voxel during the *j*th repetition (scanning run) of the *i*th word. The stability score for this voxel was then computed as the averaged pairwise correlations over all pairs of columns in the matrix (scanning runs). This produced a stability grey matter map for each subject. These stability maps were then submitted to a one-sample t-test at the group level and the voxels with top *t* values (ranging from top 100 to top 5000) were considered to be consistently high stability across subjects.

Third, we searched the term “word” in Neurosynth ^29^, which carried out an automated meta-analysis of 944 studies and produced an association test map thresholded at False discovery rate (FDR) of 0.01. Clusters with voxel sizes smaller than 10 voxels were further removed.

Fourth, in case any regions sensitive to word emotional meanings were not included, we redefined the word-associated mask as those clusters sensitive to any differences among “object vs. emotional non-object vs. non-emotional non-object”. Similar to the object vs. non-object contrast, a GLM was built to include regressors corresponding to the onsets of each of the three word types for each run. The beta maps for each word type versus baseline were submitted to a one-way ANOVA (within subject) at the group level.

Finally, instead of extracting activation patterns from a group-defined word-associated mask, we localized word-associated voxels in individual subjects using a group-constrained subject-specific approach ^30^. Adopting a leave-one-subject-pair-out procedure, we first localized group-level word-associated parcels in 19 subjects based on the object vs. non-object contrast. Within these parcels, we identified, for each of the remaining two subjects, the set of voxels showing largest differences between object and non-object words. Results of ISC-brain were largely similar when the number of individual-defined voxels increased from top 50 to 400 voxels and to all the voxels in the group-defined mask. We then united the two sets of voxels in the two subjects, and calculated Pearson correlations of activation patterns for this subject-pair for each word. For a given word, the correlations across all subject pairs were Fisher-Z-transformed and averaged to obtain the ISC-brain.

#### Brain visualization

The brain maps and results were projected onto the MNI brain surface using the BrainNet Viewer ^32^ (https://www.nitrc.org/projects/bnv/) with the default “interpolated” mapping algorithm, unless stated explicitly otherwise.

### Ratings of candidate organizing principles of semantic representations in the brain

To explain the cognitive origins of word meaning variation across individuals, we collected ratings on the following dimensions relevant to semantic representations. Each dimension was rated on a 1-7 scale, where for emotional valence 1 = negative, 4 = neutral, 7 = positive and for other ratings 1 = lowest extent, 7 = highest extent. The rating instructions were as follows:

*Sensory experience*: “to what extent the concept denoted by the word evokes a sensory experience (including vision, audition, taste, touch, and smell).”

*Navigation*: “to what extent the concept denoted by the word could offer spatial information to help you explore the environment.”

*Manipulation*: “to what extent the concept denoted by the word can be grasped easily and used with one hand.”

*Stress-related actions*: “to what extent the concept denoted by the word would make you to show a stress response, e.g., run away, attack, or freeze.”

*Emotional valence*: “to what extent the concept denoted by the word evokes positive or negative feelings: very positive feelings mean that you are happy, satisfied, contented, hopeful; very negative feelings mean that you are unhappy, annoyed, unsatisfied, despaired, or bored.”

*Arousal*: “to what extent the concept denoted by the word make you feel aroused. Low arousal means that you feel completely relaxed, very calm, sluggish, dull, or sleepy; high arousal means that you are stimulated, excited, frenzied, jittery, or wide-awake.”

*Language descriptivity*: “to what extent the concept denoted by the word could be described and explained using language.”

We recruited independent groups of 28-31 college students from Beijing Normal University for each rating via an online survey (www.sojump.com). We computed a quality metric by correlating each subject’s ratings with the averaged ratings of all but this subjects’ data across all rated words for each rating. Subjects whose ratings with others’ mean ratings were not significantly correlated (*p* > .05, uncorrected) were excluded from the subsequent analyses, leaving 24-28 college students in each rating.

### Data and materials availability

The raw data and computer code necessary to replicate the findings of the study are open to share upon request.

## Acknowledgments

We thank Bijun Wang for assistance with data collection.

## Funding

The National Natural Science Foundation of China (31671128 to Y.B, 31700943 to X.W.), Changjiang Scholar Professorship Award (T2016031 to Y.B.); the Fundamental Research Funds for the Central Universities (2017EYT35 to Y.B.), the Interdisciplinary Research Funds of Beijing Normal University (to Y.B.), and China Postdoctoral Science Foundation (2017M610791 to X.W.).

## Author contributions

XW and YB conceived and designed research; XW performed research; XW analyzed data; YB and XW wrote the paper.

## Competing interests

The authors declare no competing interests.

## Notes

### Competing Interest Statement

The authors have declared no competing interest.

