## Supplementary materials for "Idiosyncratic Tower of Babel: Individual differences in word meaning representation increase along abstractness"

**This file includes:**

Supplementary Results
Figures S1-S6
Tables S1-S5

**Supplementary results**

*PCA of semantic dimensions and their relationships with ISC-behavior and ISC-brain of word meanings*

For dimension reduction, principal component analyses (PCA) were performed on the rating means and variations (SDs), respectively. Both sets of variables were suited for PCA as suggested by the Kaiser-Meyer-Olkin measure of sampling adequacy (rating mean: 0.612; rating SD: 0.796) and Bartlett's Test of Sphericity (rating mean: chi-square (21) = 542.3, $p < .001$; rating SD: chi-square (21) = 295.1, $p < .001$). **Table S3** shows the loadings of PCs with eigenvalues above one after varimax rotation. For the 2 PCs of rating means (explaining 76.3% of total variance), PC1 had high loadings on language descriptivity, sensory, and manipulation and navigation actions; PC2 had high loadings on stress-related actions, arousal and valence. The 2 PCs of rating SDs (explaining 72.3% of total variance) had similar loading distributions on semantic dimensions with rating mean PCs.

Multiple linear regression models to predict ISC-behavior and ISC-brain using these semantic PCs are shown in **Table S4**. The magnitude of ISC-behavior could be highly significantly predicted by Mean PC1 (Language/Sensory/Navigation/Manipulation) and SD PC2 (Valence/Arousal/Stress-action): ISC-behavior = 0.78 × Mean PC1 - 0.23 × SD PC2 + 0.35 (stepwise regression model; $r^2 = .71$, $F(2,87) = 106.73$, $p = 3.85 \times 10^{-24}$). When psycholinguistic variables (i.e., word frequency and familiarity) were further included, the effects of the semantic PCs are still highly significant. The magnitude of ISD-brain could also be significantly predicted by Mean PC1 and SD PC2: ISC-brain = 0.63 × Mean PC1 + 0.18 × SD PC2 + 0.033 (stepwise regression model, $r^2 = .40$, $F(2,87) = 28.80$, $p = 2.52 \times 10^{-10}$). Note that here the SD PC2 is in the opposite direction to that in the ISC-behavior stepwise model above. When including psycholinguistic variables (i.e., word frequency, familiarity, number of strokes) into the model, the Mean PC1 becomes the only significant predictor.

Note that in regression models including psycholinguistic confounding variables, the SD PC1 was not included as an independent regressor for the following reasons. First, it was highly correlated with Mean PC1 (Pearson $r = -.87$, $p = 2.62 \times 10^{-29}$), which exacerbated multicollinearity in these regression models by increasing the variance inflation factor (VIF) of Mean PC1 to greater than 10. Second, when Mean PC1 was controlled for using partial correlation, SD PC1 became uncorrelated with ISC-behavior (Pearson $r = .006$, $p = .95$) and ISC-brain (Pearson $r = .014$, $p = .90$).

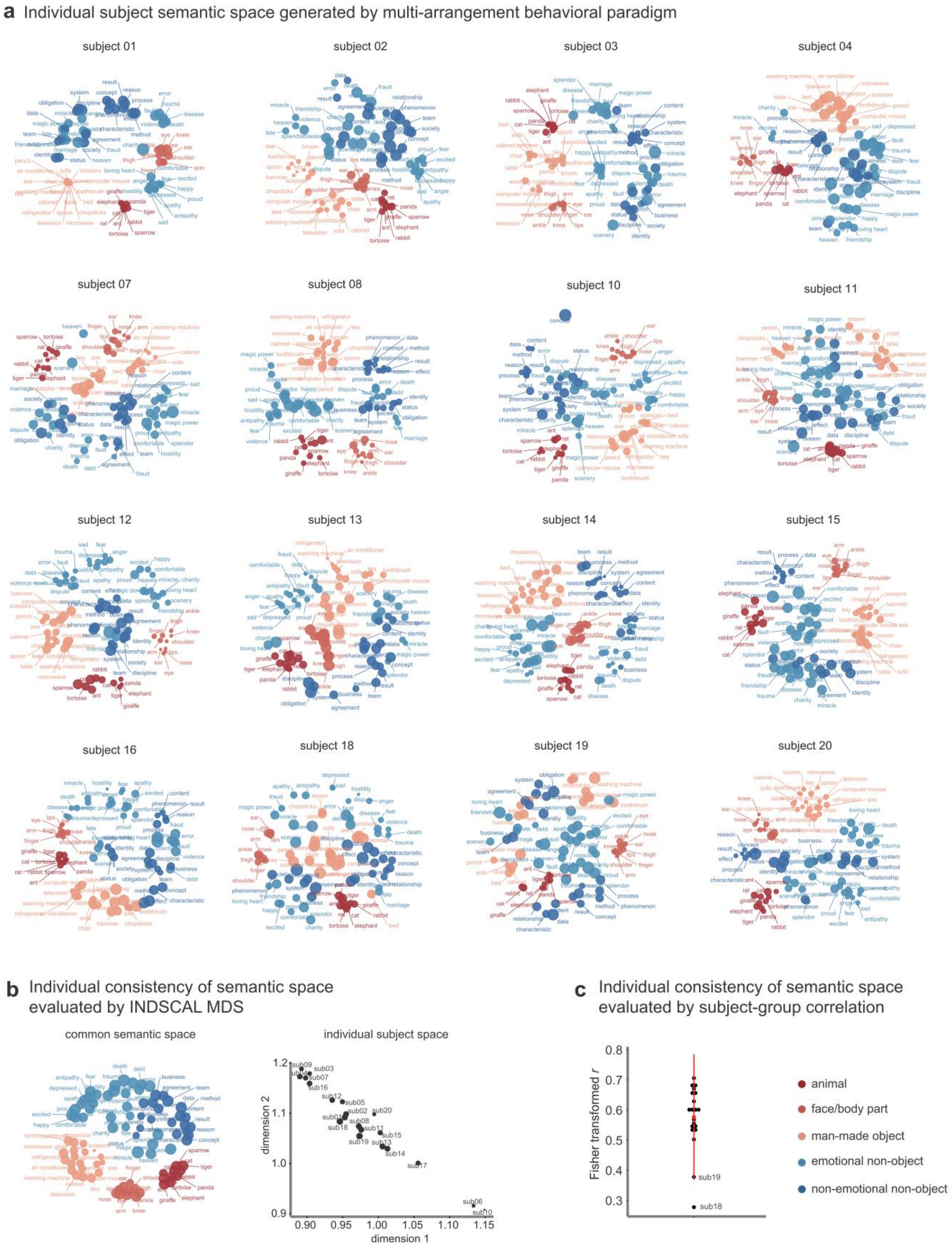
a Individual subject semantic space generated by multi-arrangement behavioral paradigm
subject 01
subject 02
subject 03
subject 04
subject 07
subject 08
subject 10
subject 11
subject 12
subject 13
subject 14
subject 15
subject 16
subject 18
subject 19
subject 20
b Individual consistency of semantic space evaluated by INDSCAL MDS
common semantic space
individual subject space
dimension 1
dimension 2
c Individual consistency of semantic space evaluated by subject-group correlation
Fisher transformed r
sub19
sub18
animal
face/body part
man-made object
emotional non-object
non-emotional non-object

**Fig. S1. (a)** Multidimensional scaling visualization of individual semantic space generated from multi-arrangement behavioral paradigm. MDS was performed using the smacof package in R (type = interval); the stress for each subject was low and indicated fair levels of fit (mean ± SD = 0.16 ± 0.05, range: 0.09-0.28). **(b)** Individual consistency of semantic space evaluated by INDSCAL MDS. The individual subject space on the right panel shows each individual subject's weights along the 2 dimensions of the common semantic space shown on the left panel. Dot sizes in above figures reflect stress, with larger dots indicating higher stress (i.e., greater inconsistency between the original and the MDS distances). **(c)** Individual consistency of semantic space evaluated by the correlations between each subject's semantic space and the leave-one-subject-out group-averaged semantic space: each black dot represents a subject, the red dot indicates group mean, and the red line indicates 1 standard deviation.

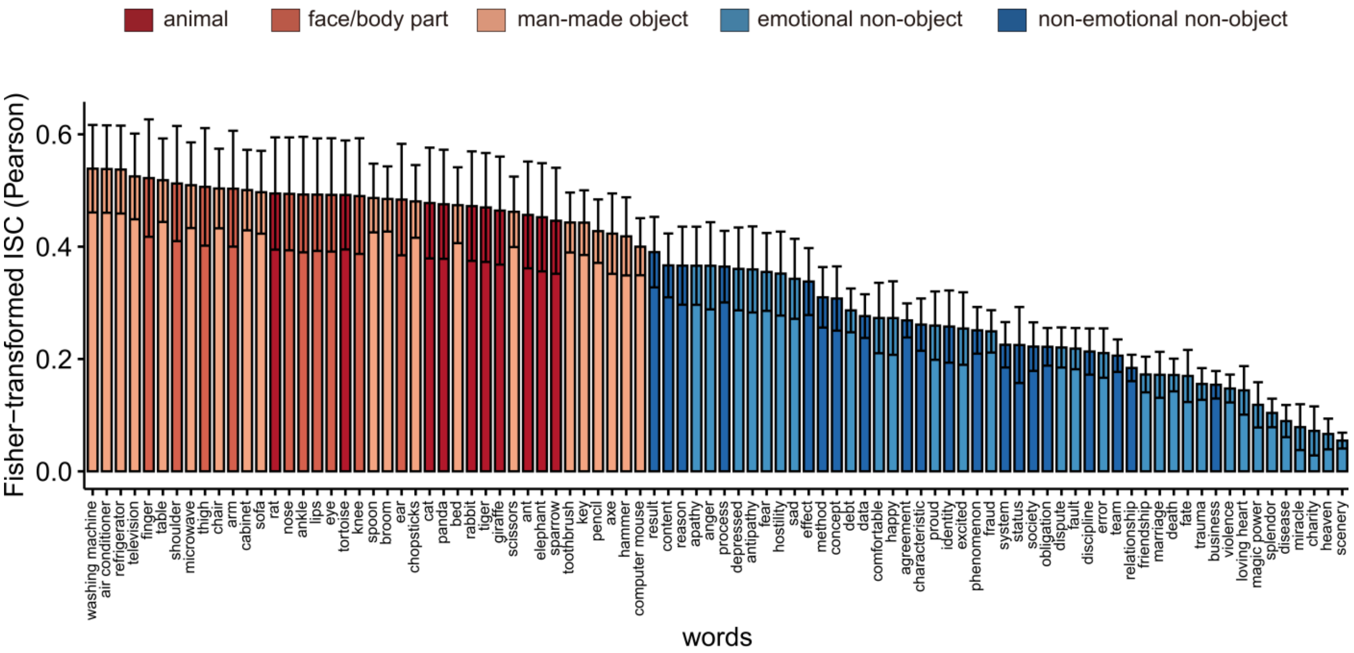


**Fig. S2.** The ISC-behavior of 90 words, in which the error bars of ISCs for each word were calculated as the standard deviations of the bootstrap distribution of 10,000 resamplings of words.

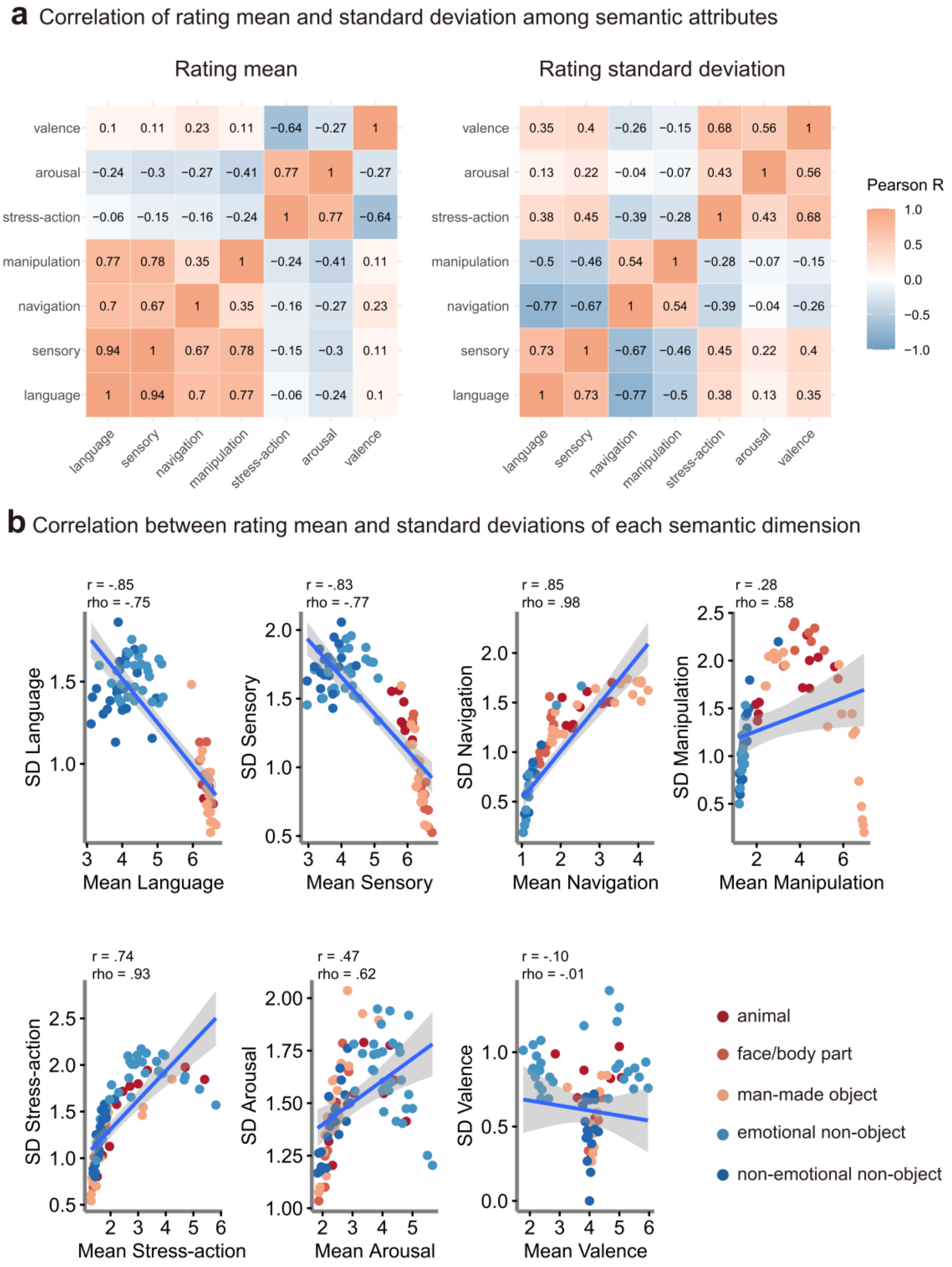


**Fig. S3. (a)** The correlations among seven semantic dimensions across 90 words in rating mean and standard deviations. **(b)** Scatter plots and correlation coefficients (Pearson R and Spearman Rho) between rating mean and SD for each semantic dimension.

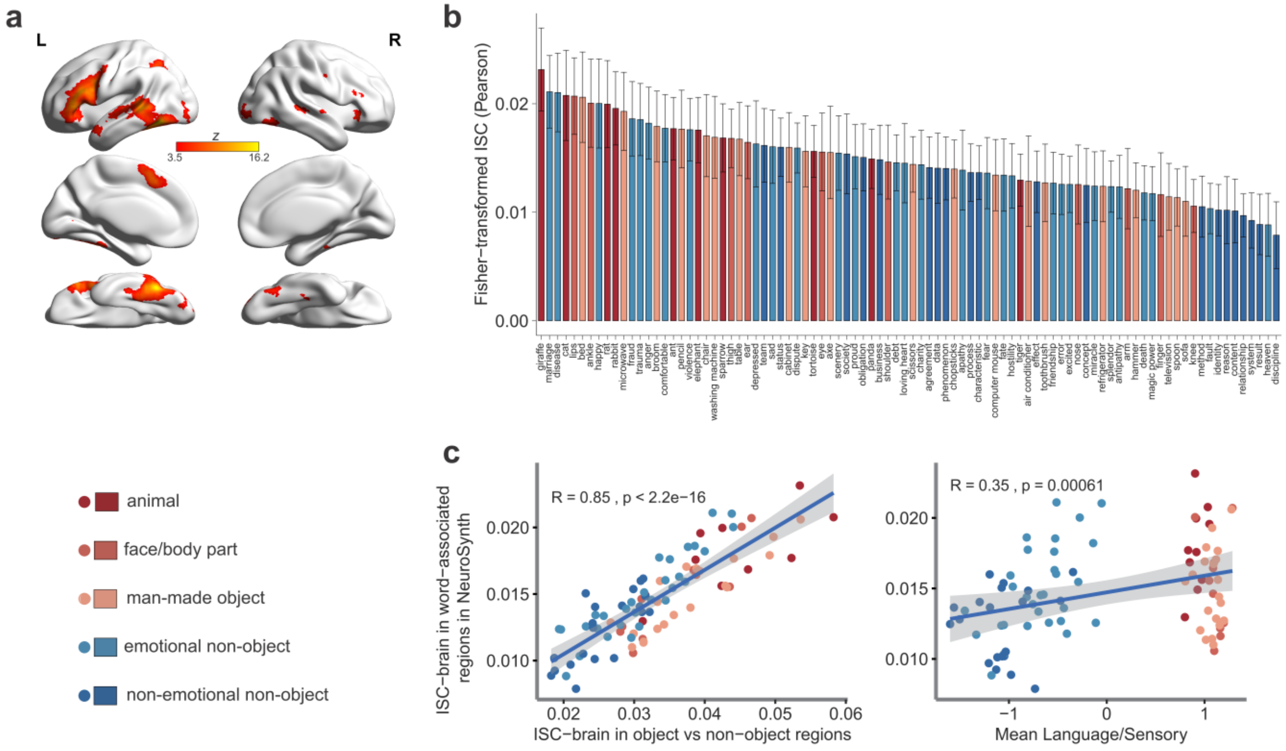


**Fig. S4.** ISC-brain calculated in Neurosynth-defined word-associated brain regions. (**a**) The meta-analytic map of 944 studies using the search term "word" in Neurosynth, which was thresholded at FDR criterion of .01. Clusters with voxel sizes smaller than 10 voxels were further removed for ISC-brain calculation. ISC-brain of 90 words in this mask (shown in **b**) was highly correlated with ISC-brain values based on the mask defined by the object vs. non-object contrast and significantly correlated with Mean Language/sensory ratings (shown in **c**).

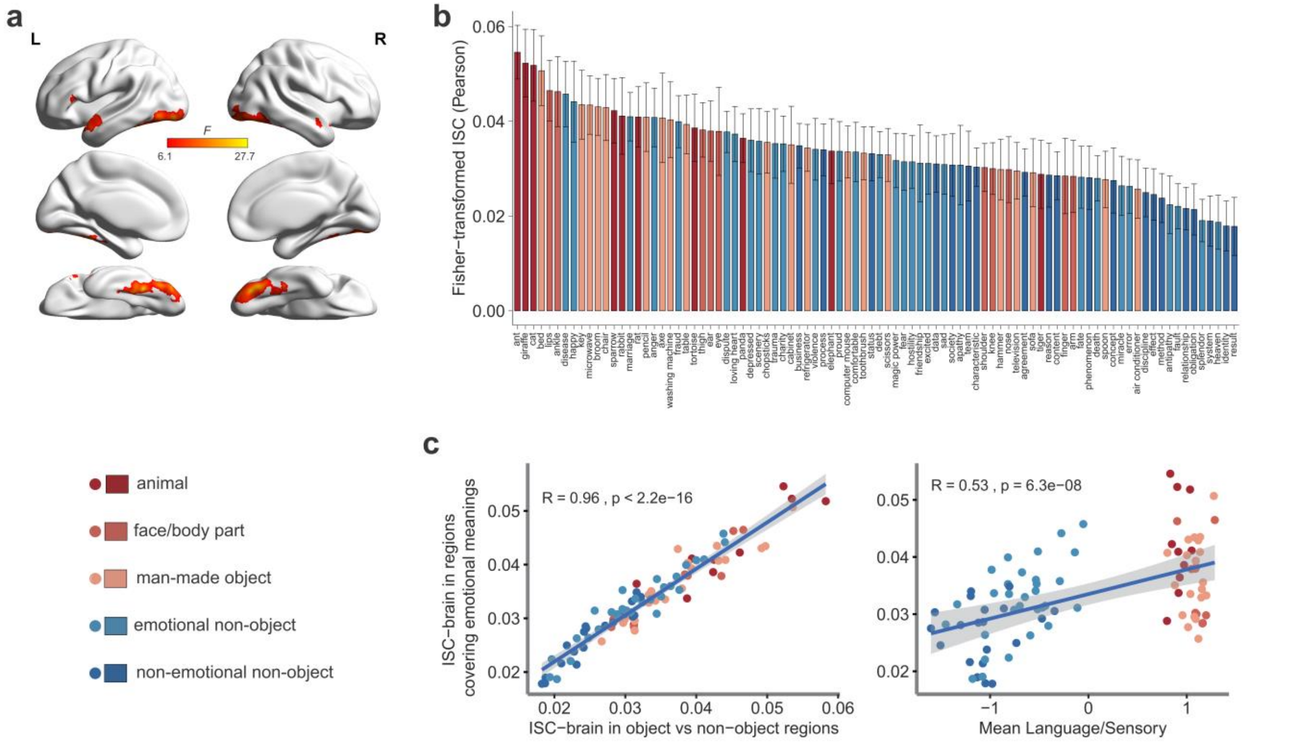


**Fig. S5.** ISC-brain calculated in brain regions defined to include regions sensitive to word emotional meanings. (**a**) These brain regions were defined as showing different activations to objects vs. emotional non-object words vs. non-emotional non-object words. The map shows F values of a one-way ANOVA (within-subject) at the group-level analysis, thresholded at voxelwise $p < .005$, FWE-corrected cluster-level $p < .05$. ISC-brain of 90 words in this mask (shown in **b**) was highly correlated with ISC-brain values based on the mask defined by the object vs. non-object contrast and significantly correlated with Mean Language/sensory ratings (shown in **c**).

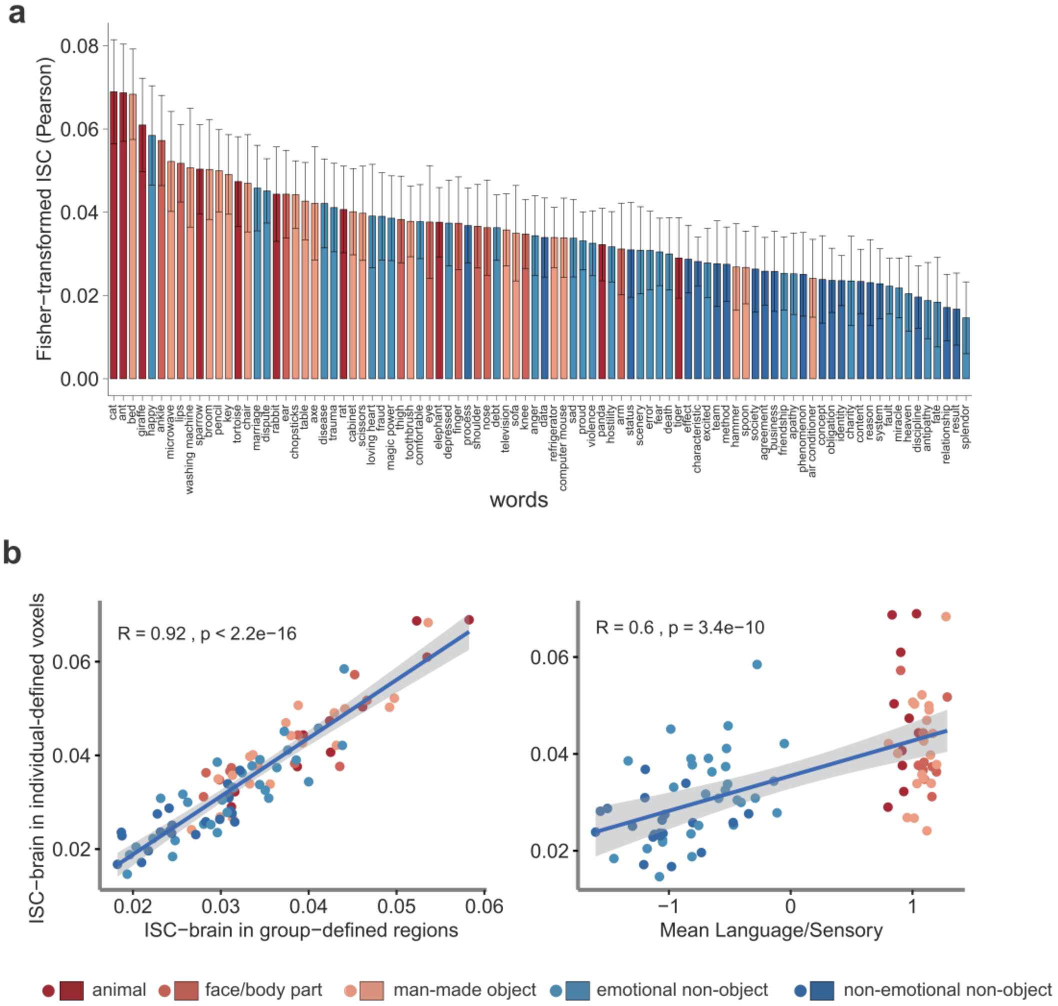


**Fig. S6.** ISC-brain calculated in individual-defined word-associated brain regions. In a leave-one-subject-pair-out procedure, meaning-related voxels were defined, in each of the two left-out subjects, as the 300 voxels showing largest differences between object and non-object words in the group mask identified in the remaining 19 subjects (F test at the group-level, voxelwise $p$ < .005, FWE-corrected cluster-level $p$ < .05). We then concatenated the 2 sets of voxels in these two subjects, extracted word $t$-values in these voxels, and computed Pearson correlations for all the words in this subject-pair. For each word, correlations across all subject pairs were Fisher-Z-transformed and averaged to obtain ISC-brain (shown in **a**). ISC-brain of 90 words in such a mask was highly correlated with ISC-brain in the main results and significantly correlated with Mean Language/sensory ratings (shown in **b**). Note that results of ISC-brain were largely similar when the number of individual-defined voxels varies from 50 to 400 and to all the voxels in the group-defined mask.

**Table S1.** Words used in this study (Chinese words and English translations)

| | Stimuli |
|---|---|
| **Words with external referents (40)** | **Animals (10):**<br>熊猫(panda); 老鼠 (rat); 老虎 (tiger); 大象 (elephant); 麻雀(sparrow); 乌龟(tortoise); 蚂蚁 (ant); 兔子 (rabbit); 猫 (cat); 长颈鹿 (giraffe);<br>**Face/body parts (10):**<br>肩膀 (shoulder); 胳膊 (arm); 大腿 (thigh); 鼻子 (nose); 眼睛 (eye); 嘴唇 (lips); 耳朵 (ear); 膝盖 (knee); 手指(finger); 脚踝(ankle);<br>**Man-made objects (20):**<br>柜子(cabinet); 椅子(chair); 沙发(sofa); 空调(air conditioner); 冰箱 (refrigerator); 电视 (television); 桌子 (table); 床 (bed); 微波炉 (microwave); 洗衣机 (washing machine); 勺子 (spoon); 剪刀 (scissors); 斧头 (axe); 鼠标 (computer mouse); 筷子 (chopsticks); 扫帚 (broom); 牙刷 (toothbrush); 钥匙 (key); 铅笔 (pencil); 锤子 (hammer) |
| **Words without external referents (50)** | **Emotional non-object words (30):**<br>爱心(loving heart); 慈善(charity);风景(scenery); 光彩(splendor); 婚姻(marriage); 骄傲(proud); 快乐(happy); 魔力(magic power);奇迹(miracle); 舒心(comfortable); 天堂(heaven); 兴奋 (excited); 缘分(fate);友情(friendship); 暴力(violence); 创伤(trauma); 错误(error); 敌意 (hostility); 反感(antipathy); 愤怒(anger); 过失(fault); 疾病(disease); 纠纷(dispute); 沮丧 (depressed); 恐惧(fear); 冷漠(apathy); 难过(sad); 骗局(fraud); 死亡(death); 债务(debt);<br>**Non-emotional non-object words (20):**<br>地位(status); 方法(method); 概念(concept); 关系(relationship); 过程(process); 纪律 (discipline); 结果(result); 买卖(business); 内容(content); 社会(society); 身份(identity); 数据 (data); 团队(team); 现象(phenomenon); 协议(agreement); 性质(characteristic); 义务 (obligation); 原因(reason); 制度(system); 作用(effect) |

**Table S2.** Multiple linear regression models to predict ISC-behavior using rating means and standard deviations of semantic dimensions

| | Unstandardized Coefficients | | | | | Variance |
|---|---|---|---|---|---|---|
| | | | Standardized | | | Inflation |
| | B | Std. Error | Coefficients | t | Sig. | Factor |
| *Predicting ISC-behavior using rating means of semantic dimensions ($R^2$ = 0.772; adjusted $R^2$ = 0.756)* | | | | | | |
| (Constant) | .635 | .073 | | 8.738 | .000 | |
| Mean Navigation | .012 | .014 | .075 | .858 | .393 | 2.775 |
| Mean Manipulation | -.008 | .008 | -.107 | -1.035 | .304 | 3.918 |
| Mean Stress-action | -.010 | .016 | -.073 | -.600 | .550 | 5.440 |
| **Mean Arousal** | **-.042** | **.015** | **-.297** | **-2.829** | **.006** | **4.008** |
| **Mean Valence** | **-.034** | **.013** | **-.220** | **-2.726** | **.008** | **2.377** |
| **Mean Language/Sensory** | **.113** | **.018** | **.780** | **6.102** | **.000** | **5.946** |
| *Predicting ISC-behavior using rating SDs of semantic dimensions ($R^2$ = 0.683; adjusted $R^2$ = 0.656)* | | | | | | |
| (Constant) | .683 | .098 | | 6.988 | .000 | |
| **SD Language** | **-.222** | **.043** | **-.574** | **-5.186** | **.000** | **3.176** |
| SD Sensory | -.027 | .035 | -.076 | -.769 | .444 | 2.529 |
| SD Navigation | -.003 | .029 | -.011 | -.107 | .915 | 2.893 |
| **SD Manipulation** | **.043** | **.019** | **.170** | **2.260** | **.026** | **1.469** |
| SD Stress-action | .011 | .028 | .034 | .382 | .703 | 2.101 |
| SD Arousal | -.026 | .048 | -.041 | -.539 | .591 | 1.532 |
| **SD Valence** | **-.103** | **.043** | **-.229** | **-2.422** | **.018** | **2.321** |
| *Predicting ISC-behavior using significant rating means and SDs of semantic dimensions ($R^2$ = 0.772; adjusted $R^2$ = 0.756)* | | | | | | |
| (Constant) | .629 | .068 | | 9.224 | .000 | |
| **Mean Language/Sensory** | **.080** | **.017** | **.554** | **4.813** | **.000** | **4.827** |
| **Mean Arousal** | **-.030** | **.014** | **-.217** | **-2.219** | **.029** | **3.479** |
| **Mean Valence** | **-.027** | **.009** | **-.173** | **-3.078** | **.003** | **1.147** |
| SD Language | -.067 | .043 | -.175 | -1.565 | .121 | 4.542 |
| SD Manipulation | .018 | .017 | .070 | 1.070 | .288 | 1.575 |
| SD Valence | -.042 | .041 | -.093 | -1.012 | .314 | 3.050 |
| *Predicting ISC-behavior using significant rating means and SDs of semantic dimensions and psycholinguistic confounding variables ($R^2$ = 0.795; adjusted $R^2$ = 0.775)* | | | | | | |
| (Constant) | -.108 | .255 | | -.425 | .672 | |
| **Mean Language/Sensory** | **.044** | **.021** | **.302** | **2.088** | **.040** | **8.287** |
| **Mean Arousal** | **-.032** | **.014** | **-.225** | **-2.323** | **.023** | **3.720** |
| **Mean Valence** | **-.033** | **.009** | **-.211** | **-3.814** | **.000** | **1.212** |
| SD Language | -.064 | .041 | -.165 | -1.541 | .127 | 4.548 |
| SD Manipulation | .023 | .016 | .090 | 1.420 | .159 | 1.603 |
| SD Valence | -.028 | .040 | -.063 | -.703 | .484 | 3.160 |
| word frequency | -.009 | .012 | -.049 | -.784 | .435 | 1.557 |
| **word familiarity** | **.117** | **.039** | **.293** | **2.978** | **.004** | **3.840** |

**Table S3.** Semantic variables for principal component analyses (PCA) and the resulting component loadings under varimax rotation

| PCA on rating means | Mean PC1 | Mean PC2 | PCA on rating standard deviations (SD) | SD PC1 | SD PC2 |
|---|---|---|---|---|---|
| Variance explained | 49.89% | 26.38% | Variance explained | 50.11% | 22.21% |
| *Loadings on each dimension* | | | *Loadings on each dimension* | | |
| Mean Language | **0.976** | -0.013 | SD Navigation | **-0.894** | -0.090 |
| Mean Sensory | **0.964** | -0.083 | SD Language | **0.875** | 0.187 |
| Mean Manipulation | **0.811** | -0.197 | SD Sensory | **0.799** | 0.311 |
| Mean Navigation | **0.739** | -0.151 | SD Manipulation | **-0.727** | -0.024 |
| Mean Stress-action | -0.054 | **0.966** | SD Valence | 0.201 | **0.871** |
| Mean Arousal | -0.274 | **0.785** | SD Arousal | -0.055 | **0.831** |
| Mean Valence | 0.035 | **-0.745** | SD Stress-action | 0.351 | **0.758** |

**Table S4.** Multiple linear regression models to predict ISC-behavior and ISC-brain using principal components extracted from rating means and standard deviations of semantic dimensions

| | Unstandardized Coefficients | | Standardized Coefficients | t | Sig. | Variance Inflation Factor |
|---|---|---|---|---|---|---|
| | B | Std. Error | | | | |
| *Predicting ISC-behavior using semantic PCs ($R^2$ = 0.715; adjusted $R^2$ = 0.702)* | | | | | | |
| (Constant) | .346 | .008 | | 42.239 | .000 | |
| **Mean PC1 (Language/Sensory/Navigation/Manipulation)** | **.104** | **.023** | **.728** | **4.442** | **.000** | **8.005** |
| Mean PC2 (Stress-action/Arousal/Valence) | -.007 | .013 | -.052 | -.578 | .565 | 2.409 |
| SD PC1 (Language/Sensory/ Navigation/Manipulation) | -.009 | .024 | -.062 | -.373 | .710 | 8.224 |
| **SD PC2 (Valence/Arousal/Stress-action)** | **-.030** | **.012** | **-.208** | **-2.428** | **.017** | **2.190** |
| *Predicting ISC-behavior using semantic PCs and psycholinguistic confounding variables ($R^2$ = 0.736; adjusted $R^2$ = 0.720) ** | | | | | | |
| (Constant) | -.293 | .258 | | -1.134 | .260 | |
| **Mean PC1 (Language/Sensory/Navigation/Manipulation)** | **.085** | **.016** | **.595** | **5.462** | **.000** | **3.771** |
| Mean PC2 (Stress-action/Arousal/Valence) | -.002 | .010 | -.017 | -.247 | .806 | 1.508 |
| **SD PC2 (Valence/Arousal/Stress-action)** | **-.027** | **.009** | **-.190** | **-2.887** | **.005** | **1.370** |
| word frequency | .001 | .013 | .008 | .116 | .908 | 1.454 |
| **word familiarity** | **.098** | **.041** | **.246** | **2.423** | **.018** | **3.291** |
| *Predicting ISC-brain using semantic PCs ($R^2$ = 0.403; adjusted $R^2$ = 0.374)* | | | | | | |
| (Constant) | .033 | .001 | | 45.503 | .000 | |
| **Mean PC1 (Language/Sensory/Navigation/Manipulation)** | **.007** | **.002** | **.790** | **3.329** | **.001** | **8.005** |
| Mean PC2 (Stress-action/Arousal/Valence) | .000 | .001 | -.036 | -.273 | .786 | 2.409 |
| SD PC1 (Language/Sensory/Navigation/Manipulation) | .002 | .002 | .172 | .717 | .475 | 8.224 |
| SD PC2 (Valence/Arousal/Stress-action) | .002 | .001 | .216 | 1.744 | .085 | 2.190 |
| *Predicting ISC-brain using semantic PCs and psycholinguistic confounding variables ($R^2$ = 0.416; adjusted $R^2$ = 0.374) ** | | | | | | |
| (Constant) | .013 | .024 | | .528 | .599 | |
| **Mean PC1 (Language/Sensory/Navigation/Manipulation)** | **.004** | **.001** | **.502** | **3.058** | **.003** | **3.833** |
| Mean PC2 (Stress-action/Arousal/Valence) | .000 | .001 | .036 | .353 | .725 | 1.508 |
| SD PC2 (Valence/Arousal/Stress-action) | .001 | .001 | .156 | 1.588 | .116 | 1.370 |
| word frequency | -.001 | .001 | -.069 | -.662 | .510 | 1.538 |
| word familiarity | .003 | .004 | .120 | .782 | .436 | 3.349 |
| number of strokes | .000 | .000 | .090 | 1.041 | .301 | 1.067 |

*: The SD PC1 was not included in these regression models due to its high correlation with Mean PC1.

**Table S5.** Multiple linear regression models to predict ISC-brain using rating means and standard deviations of semantic dimensions

| | Unstandardized Coefficients B | Unstandardized Coefficients Std. Error | Standardized Coefficients | t | Sig. | Variance Inflation Factor |
|---|---|---|---|---|---|---|
| *Predicting ISC-brain using rating means of semantic dimensions ($R^2$ = 0.404; adjusted $R^2$ = 0.361)* | | | | | | |
| (Constant) | .028 | .007 | | 3.884 | .000 | |
| **Mean Language/Sensory** | **.005** | **.002** | **.578** | **2.795** | **.006** | **5.946** |
| Mean Navigation | .000 | .001 | .047 | .332 | .741 | 2.775 |
| Mean Manipulation | .000 | .001 | .079 | .472 | .638 | 3.918 |
| Mean Stress-action | .000 | .002 | -.055 | -.279 | .781 | 5.440 |
| Mean Arousal | .002 | .001 | .226 | 1.330 | .187 | 4.008 |
| Mean Valence | .000 | .001 | -.045 | -.344 | .732 | 2.377 |
| *Predicting ISC-brain using rating SDs of semantic dimensions ($R^2$ = 0.356; adjusted $R^2$ = 0.301)* | | | | | | |
| (Constant) | .036 | .009 | | 4.139 | .000 | |
| **SD Language** | **-.012** | **.004** | **-.488** | **-3.088** | **.003** | **3.176** |
| SD Sensory | .000 | .003 | .013 | .094 | .925 | 2.529 |
| SD Navigation | .003 | .003 | .195 | 1.294 | .199 | 2.893 |
| SD Manipulation | .000 | .002 | .009 | .086 | .932 | 1.469 |
| SD Stress-action | .004 | .003 | .225 | 1.749 | .084 | 2.101 |
| SD Arousal | .002 | .004 | .041 | .376 | .708 | 1.532 |
| SD Valence | -.001 | .004 | -.021 | -.158 | .875 | 2.321 |
| *Predicting ISC-brain using significant rating means and SDs of semantic dimensions ($R^2$ = 0.376; adjusted $R^2$ = 0.361)* | | | | | | |
| (Constant) | .036 | .005 | | 7.384 | .000 | |
| SD Language | -.002 | .004 | -.083 | -.498 | .620 | 3.888 |
| **Mean Language/Sensory** | **.005** | **.001** | **.540** | **3.232** | **.002** | **3.888** |
| *Predicting ISC-brain using significant rating means and SDs of semantic dimensions and psycholinguistic variables ($R^2$ = 0.391; adjusted $R^2$ = 0.355)* | | | | | | |
| (Constant) | .033 | .026 | | 1.283 | .203 | |
| SD Language | -.003 | .004 | -.112 | -.657 | .513 | 3.982 |
| **Mean Language/Sensory** | **.004** | **.002** | **.450** | **2.012** | **.047** | **6.906** |
| word frequency | -.001 | .001 | -.107 | -1.032 | .305 | 1.478 |
| word familiarity | .000 | .004 | .019 | .120 | .905 | 3.546 |
| number of strokes | .000 | .000 | .068 | .785 | .435 | 1.050 |
